# Lateralization of feeding behaviour in white-fronted lemur (*Eulemur albifrons*) and ring-tailed lemur (*Lemur catta*) in captivity

**DOI:** 10.1101/2023.12.21.572747

**Authors:** Laura Calvo Heredia, Francisco Javier de Miguel Águeda

## Abstract

Functional cerebral asymmetry is reflected in the lateralization of some behavioural patterns in many vertebrate species. In primates, behavioural lateralization has been related to both life style and age and sex, and it affects behaviours such as feeding and other tasks that require precision movements.

We have studied feeding lateralization concerning the use of right and left hand to take the food in two species of lemurs, the mainly arboreal white-fronted lemur and the more terrestrial ring-tailed lemur, taking also account the age and the sex of the individuals. Half of the white-fronted lemurs (7 of 14) showed lateralization in feeding, while only a few ring-tailed lemurs (3 of 19) showed it. In the first species, a light bias seems to emerge (5 individuals used mostly the right hand for taking the food, while only 2 used mainly the left hand), while in the second species no bias could really be appreciated. No clear effect of age and sex on the presence and direction of lateralization could be evidenced.

The results somehow contrast with what the postural theory of lateralization postulates about the preferential use of the right hand in terrestrial species.

## Introduction

Traditionally, brain asymmetry and behavioural lateralization were thought to be unique attributes of the human species. However, in recent decades, extensive research has revealed that these traits are widespread among various vertebrate species (Laska & Tutsch, 2000; Hopkins, 2006, Corballis et al., 2012; Regaiolli et al., 2016). Lateralization, which is defined as the uneven distribution of functions in the cerebral hemispheres resulting from cerebral asymmetry (Watson & Hanbury, 2007; Batist & Mayhew, 2020), significantly shapes an individual’s behaviour (Vallortigara & Rogers, 2005; Koboroff et al., 2008). While the brain and the nervous system are symmetrically organized in terms of sensorimotor activities, they exhibit a range of systematic and functional asymmetries. These asymmetries appear to enhance brain capacity and efficiency by avoiding redundant functions (Vallortigara & Rogers, 2020; Rogers, 2021).

This phenomenon of brain specialization is well-documented across a wide range of species, from insects to mammals (Laska & Tutsch, 2000; Hopkins, 2006; Regaiolli et al., 2016; Rogers, 2021). Research involving fish, amphibians, and reptiles has played a crucial role in elucidating the origins of lateralization in vertebrates (Rogers & Andrews, 2002; Stancher et al., 2018). These studies support the overarching idea that, broadly speaking, the right hemisphere tends to respond to potent, instinctive stimuli like predator avoidance and general social interactions. In contrast, the left hemisphere primarily handles sustained responses to particular stimuli or tasks, such as foraging for food or using tools, observed in specific species (Rogers, 2021).

Furthermore, recent research conducted over the past three decades firmly establishes that the majority of vertebrates exhibit lateral biases in the use of the limbs, i.e., handedness. This phenomenon is particularly pronounced in various mammal species, with nonhuman primates attracting particular attention due to their advanced brain development and cognitive complexity, apart from their phylogenetic relation with humans (Fagot & Vauclair, 1991; Laska & Tutsch, 2000). Although no non-human primate study has revealed lateralization as pronounced as that observed in humans, most of whom typically use the right hand for motor actions that require precision (Hopkins, 1995; Hopkins, 2006; Hopkins et al., 2011), several studies have demonstrated significant biases in different species (Ward et al., 1990; Hopkins et al., 2011; De Andrade & de Sousa 2018; Caspar et al., 2018).

Most studies on strepsirrhines, for instance, have identified a preference for using the left hand in manual tasks/motor activities (Sanford & Ward, 1986; Forsythe & Ward, 1988; Masataka 1989; Milliken et al. 1989; Ward et al. 1990; Stafford et al. 1993; Bennett et al. 1995), while a greater inclination to use the right hand has been observed in hominids for manual gestures, bimanual feeding, and simple reaching (Hopkins, 2006; Meguerditchian, et al. 2010; Schnoell et al., 2014; Pointdexter et al., 2018) and capuchin monkeys and baboons for bimanual tasks (Spinozzi et al. 1998; Vauclair et al. 2005). Prosimians, despite their more “ancestral” phylogenetic position, constitute an intriguing group of interest. They hold a crucial taxonomic position in primate phylogeny, rendering them highly compelling subjects for investigating behavioural asymmetries.

Examples of lateralization, especially concerning hand preference, have been found in sifaka (*Propithecus coquereli*) (Milliken et al., 2005), aye-aye (*Daubentonia madagascariensis*) (Lhota et al., 2009), and different species of lemurids (Ward et al., 1990; Shaw et al., 2004; Papademetriou et al., 2005; Regaiolli et al., 2016), on which we precisely have focused our study.

Lemurs, as strepsirrhine primates endemic to Madagascar, inhabit a distinctive ecological niche. These primarily crepuscular and arboreal social creatures often congregate in groups of variable size (Eppley et al., 2015). Watson and Hanbury (2007) undertook a comprehensive review that consolidated existing data on lateralization in selected prosimian species. Their findings revealed that most species, including the ring-tailed lemur (*Lemur catta*) (Milliken et al., 1989) and the white-fronted lemur (*Eulemur albifrons*), predominantly use their left hand in different manual activities (Ward et al., 1990; Papademetriou et al., 2005). However, more recently Regaiolli et al. (2016) have indicated that ring-tailed lemurs exhibit a tendency to use their right foreleg for both unimanual and bimanual experimental tasks.

Several factors appear to influence lateralization in these species. It has been proposed that the direction of lateralization may depend on factors such as posture and locomotion (Hopkins et al., 2011). Additionally, the postural origin of handedness theory (MacNeilage et al., 1987) postulated that in prosimians, the left hand, governed by the right hemisphere, has specialized for unimanual predatory prehension, while the right hand, controlled by the left hemisphere, has specialized for grasping and supporting on branches. The tendency to employ the right hand for precision tasks may have evolved in terrestrial primates, which no longer rely on it for bracing on branches.

Furthermore, individual characteristics such as age and sex can also impact lateralization in prosimians. For instance, among ring-tailed lemurs, more pronounced manual lateralization has been observed in adults compared to juveniles, a pattern observed in various species within the genus *Eulemur* as well, according to Ward et al. (1990). These authors also noted that the use of the left hand within the genus *Lemur* is more prevalent in males and juveniles.

Since the results mentioned above are somehow fragmentary, our study seeks to shed additional light on this topic by investigating lateralization in feeding behaviour among white-fronted lemurs and ring-tailed lemurs, as this behaviour has been a focal point in previous research on lateralization in prosimians (Masataka, 1989; Milliken et al., 1989; Ward et al., 1990; Milliken et al., 1991). These two species exhibit differences in various traits, including displacement between trees; the first is more arboreal, while the latter is more terrestrial. Our study aims to uncover distinctions between these species that could arise from their distinct lifestyles, and/or potentially from differences in age or sex.

## Methods

### Enclosure and subjects

White-fronted lemurs and ring-tailed lemurs share an enclosure at Biological Park Faunia (Madrid, Spain) with red ruffed lemurs *Varecia rubra* (included in a specific breeding program, thus they did not remain in this enclosure during the whole study period) and several birds species. The enclosure, in the “African Forest” (Fig. 1), has more than 1800 m^2^, is almost circular and it is closed and limited by a mesh at the top. It disposes of two entrances for visitors, keepers and park staff, and contains diverse trees and shrubs species, as well as several elements of environmental enrichment (fallen logs, artificial grass, three wooden porches and a small pond with a waterfall), and two heated shelters. The entire facility is crossed by a trail for the use of visitors, who are forbidden from interacting with the animals. The group of white-fronted lemurs was made up of 14 individuals, 7 males and 7 females (1 born in 2019 and 3 born in the spring of 2018, the rest being adults), while the group of ring-tailed lemurs consisted of 37 individuals. All white-fronted lemurs (Table 1) and 19 ring-tailed lemurs (Table 2) were included in the study.

**Figure 1.**
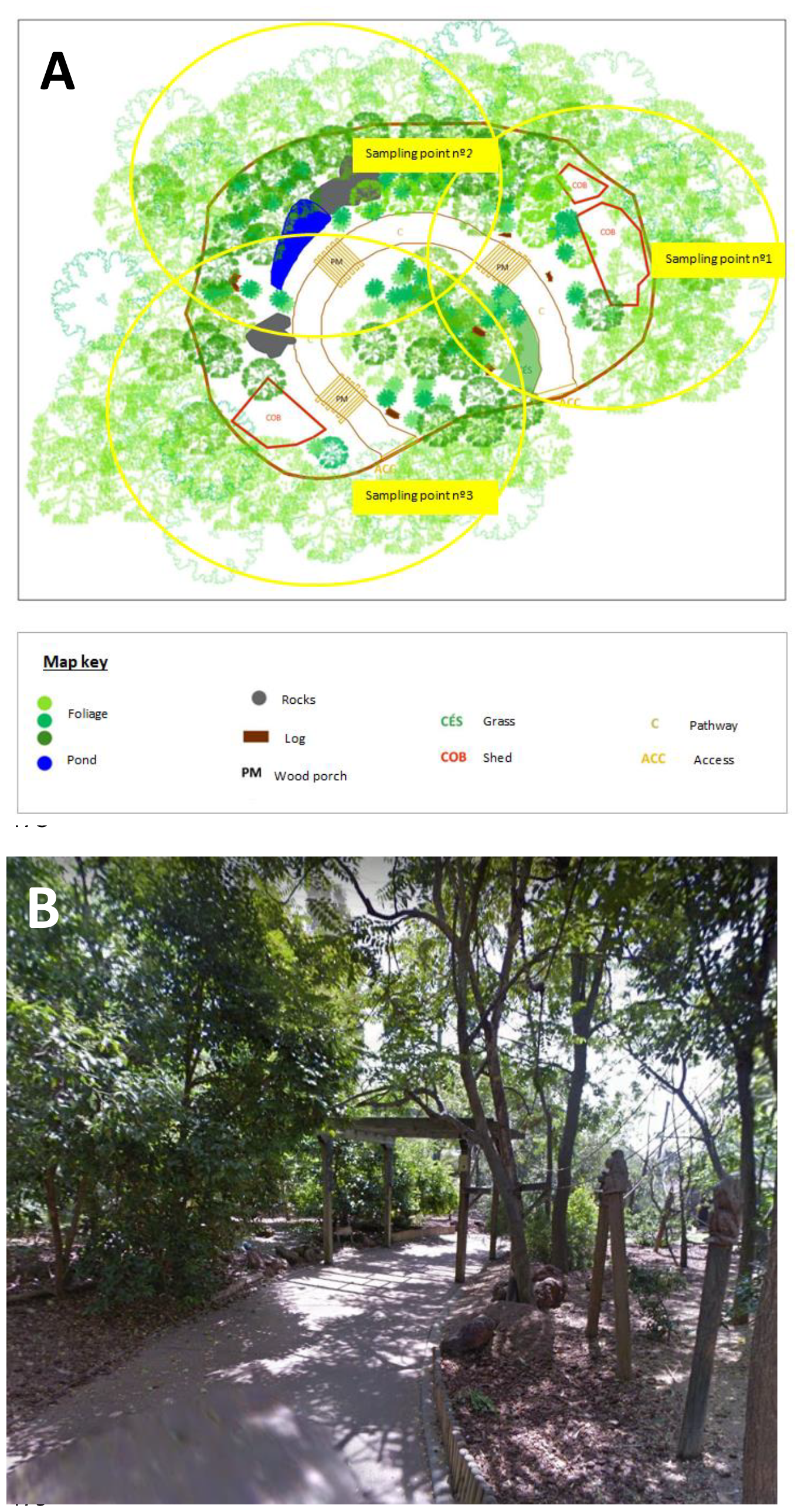
Enclosure of lemurs in Faunia: A) general view with the sampling areas indicated; B) path, vegetation and elements of environmental enrichment in the enclosure. Images obtained from The Land Registry and Google Maps

**Table 1.** White-fronted lemurs. Relationship of the studied individuals.

| Name | Code Number | Sex | Birth year | Descendant of |
| --- | --- | --- | --- | --- |
| Barouk | 1 | Male | 2004 | - |
| Bony | 2 | Male | 2001 | - |
| Reny´s baby | 3 | Female | 2019 | Reny |
| Riplie´s baby | 4 | Male | 2019 | Riplie |
| Runa´s baby | 5 | Female | 2019 | Runa |
| Lemming | 6 | Male | 2007 | - |
| Loa | 7 | Female | 2012 | Reny |
| Lui | 8 | Male | 2010 | - |
| Padre Barouk | 9 | Male | 2000 | - |
| Reny | 10 | Female | 2010 | - |
| Riplie | 11 | Female | 2016 | - |
| Riñonera | 12 | Female | 2018 | Reny |
| Rolo | 13 | Male | 2017 | Reny |
| Runa | 14 | Female | 2017 | Reny |

**Table 2.** Ring-tailed lemurs. Relationship of the studied individuals.

| Name | Code Number | Sex | Birth year | Descendant of |
| --- | --- | --- | --- | --- |
| Efea's baby | 1 | Male | 2019 | Efea |
| Ovi's baby | 2 | Male | 2019 | Ovi |
| Carapan | 3 | Female | 2016 | Ojos de huevo |
| Vladimira | 4 | Female | 2019 | Vandi |
| Ed | 5 | Female | 2001 | - |
| Efea | 6 | Female | 2016 | Ed |
| Elora | 7 | Female | 2018 | Efatra |
| Efatra | 8 | Female | 2013 | Ed |
| Igor | 9 | Male | 2004 | - |
| Kite | 10 | Male | 2001 | - |
| Ojos de huevo | 11 | Female | 2001 | - |
| Ohana | 12 | Female | 2018 | Ojos de huevo |
| Ovi | 13 | Female | 2015 | Ojos de huevo |
| Panoli | 14 | Male | 2004 | - |
| Pinocha | 15 | Female | 2003 | - |
| Pinocho | 16 | Male | 2018 | Pinocha |
| Seigi | 17 | Male | 2003 | - |
| Vampi | 18 | Female | 2002 | - |
| Vandi | 19 | Female | 2015 | Vampi |

### Procedure

The study was conducted between February and July 2019, 2-4 days a week. The first weeks were spent in the recognition of individuals by means of *ad libitum* sampling. The feeding behaviour we recorded was the concerning to the use of right and left hand by lemurs when taking the food. Data were recorded by the same observer in two time slots, 10:00-12:30 and 15:00-17:30. Behavioural sampling with continuous recording was used from three different points, in each of which the observer stayed for 50 minutes.

Sex and age of the individuals were also recorded (Tables 1 and 2). According to age, individuals were classified as adults (sexually mature individuals), juveniles (individuals born in 2018) and infants (individuals born in 2019).

### Data analysis

According to Hopkins (1999) we calculated for each individual the hand preference (if it was the case) and the strength of such preference. For each subject, a z score based on the total frequency of right-hand and left-hand responses was determined according to the next formula:

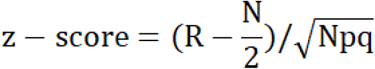

where R is the number of right hand preferences, N the total number of counts, and p = q = 0.5. Individuals with Z-scores > 1.96 were considered rightward lateralized, those having Z-scores < -1.96 were considered leftward lateralized, and animals having Z-scores between -1.96 and 1.96 were considered ambilateral.

To determine the strength of laterality we calculated the handedness index (HI), in the following way:

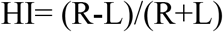

Here R is the number of right lateral biased responses, and L is the number of left lateral biased responses. Negative values represent left lateral biases, and positive values represent right lateral biases, while the absolute value of each dominance index reflects the strength of the bias.

Finally, to determine for each species the existence of right/left bias at group level, we analysed the corresponding z-scores with t-Student test.

## Results

In white-fronted lemur, 8 individuals showed a clear lateralization (Table 3), of which 2 were left-biased and 6 were right-biased. All the individuals were adult but one, a juvenile female right-biased. Among the lateralized adults, 2 males were left-biased, and 3 males and 2 females were right-biased. Individuals 7 (adult female), 12 (juvenile female) and 13 (adult male) are descendant of 10 (adult female). All of them were right-biased. Concerning the strength of lateralization, only one individual, the adult female 10, was strongly right biased, since HI= 1. Other individual, the infant female 3, had also HI=1, but she only provided 2 feeding behaviour records. No bias was found at group level (t= 1.25; p= 0.23).

**Table 3.**
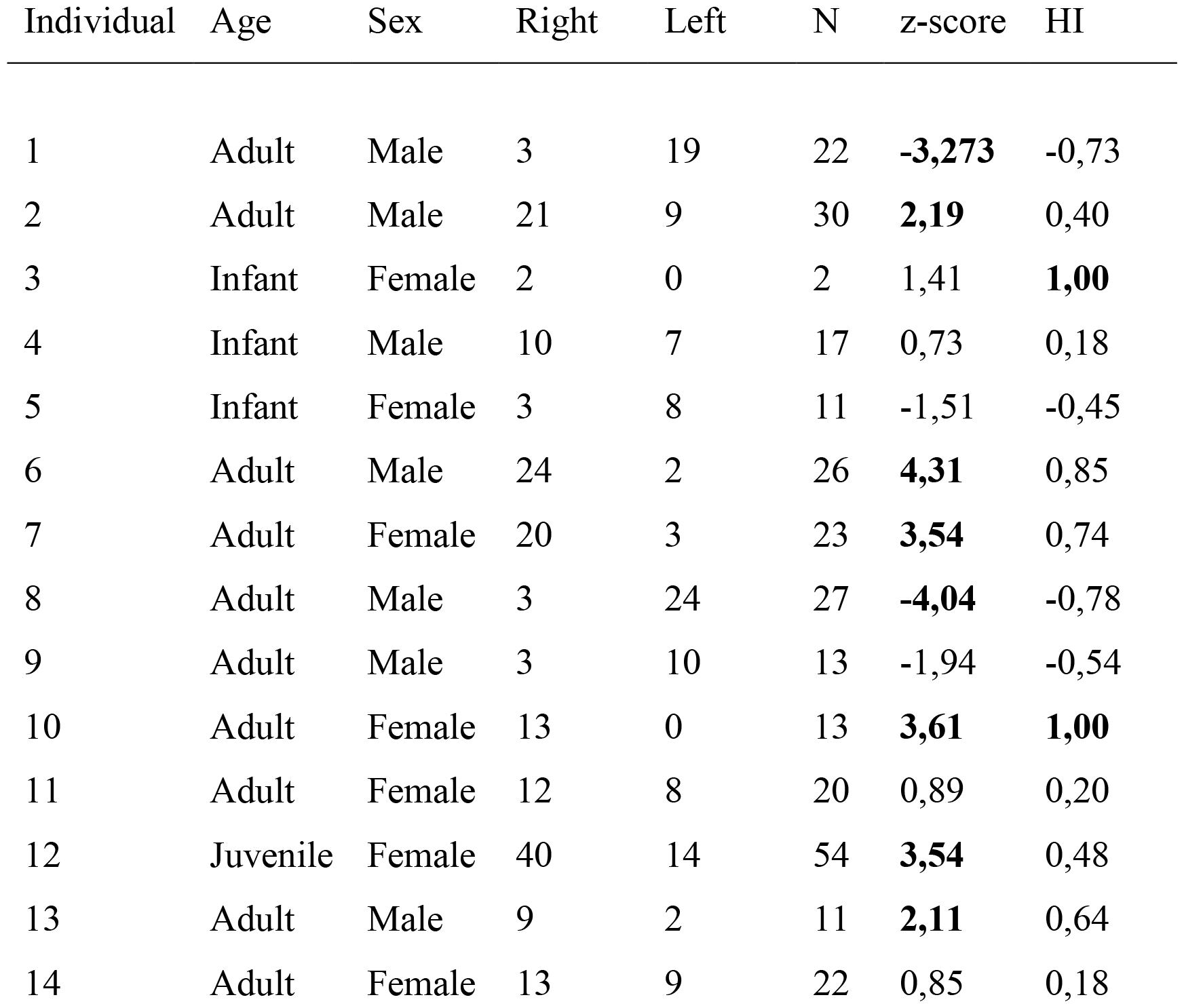
Hand preference for feeding on white-fronted lemur. Values in bold for z-score indicate significant laterality, while values in bold for HI indicate strong laterality.

With respect to ring-tailed lemur, only 3 adult individuals showed significant differences (Table 4): a male left-biased, a male right-biased and a female right-biased. No individual showed strong lateralization, either rightward or leftward. No bias was found at group level in this case too (t= 0.25; p= 0.80).

**Table 4.** Results of binomial tests on ring-tailed lemur. Values in bold for z-score indicate significant laterality, while values in bold for HI indicate strong laterality.

| Individual | Age | Sex | R | L | z-score | HI |
| --- | --- | --- | --- | --- | --- | --- |
| 1 | Infant | Male | 13 | 14 | -0,19 | -0,04 |
| 2 | Infant | Male | 4 | 10 | -1,60 | -0,43 |
| 3 | Adult | Female | 1 | 2 | -0,58 | -0,33 |
| 4 | Infant | Female | 13 | 11 | 0,41 | 0,08 |
| 5 | Adult | Female | 18 | 10 | 1,51 | 0,29 |
| 6 | Adult | Female | 4 | 10 | -1,60 | -0,43 |
| 7 | Juvenile | Female | 10 | 18 | -1,51 | -0,29 |
| 8 | Adult | Female | 10 | 7 | 0,73 | 0,18 |
| 9 | Adult | Male | 9 | 5 | 1,07 | 0,29 |
| 10 | Adult | Male | 3 | 13 | **-3,54** | -0,63 |
| 11 | Adult | Female | 15 | 1 | **3,50** | 0,88 |
| 12 | Juvenile | Female | 17 | 28 | -1,64 | -0,24 |
| 13 | Adult | Female | 0 | 0 | 0,00 | 0 |
| 14 | Adult | Male | 17 | 8 | 1,80 | 0,36 |
| 15 | Adult | Female | 10 | 3 | 1,94 | 0,54 |
| 16 | Juvenile | Male | 26 | 14 | 1,90 | 0,30 |
| 17 | Adult | Male | 15 | 6 | **1,96** | 0,43 |
| 18 | Adult | Female | 3 | 8 | -1,51 | -0,45 |
| 19 | Adult | Female | 8 | 11 | -0,69 | -0,16 |

## Discussion

In general terms, our results seem to suggest that feeding behaviour in white-fronted lemurs is more lateralized than in ring-tailed lemur. Besides, at least as far as our group is concerned, white-fronted lemurs were mostly right-handed, although only one individual showed a strong right-bias, and no bias was found at group level. On the contrary, no trend could be evidenced in ring-tailed lemur, since only three individuals showed lateralization. None of these individuals showed a strong lateralization and no bias was found at level group too.

The posture adopted when performing an activity would be a determining factor as to whether the animals will show a tendency to use either side of the body or right or left limbs (Shaw et al., 2004; Hopkins et al., 2011). According to the postural origin of handedness (MacNeilage et al., 1987), it would expected that ring-tailed lemur, more terrestrial, depended more on right hand for precision tasks like feeding, since it would no longer have to depend on this hand for grip and support on the branches. Our results have not confirmed this assumption, given that it was precisely in white-fronted lemur, more arboreal, in which a light trend to use the right hand seems to evidence. This results also contrast with those obtained by Ward et al. (1990) and Papademetriou et al. (2005) relative to dominance of left hand in manual tasks. Notwithstanding, results from other studies on lemurs, like of the Scheumann et al. (2011) and Schnoell et al. (2014), have no supported the postural origin of handedness.

However, the small sample sizes of both species in our study makes it necessary to take these results with caution, since only 2 white-fronted lemurs and 2 ring-tailed lemurs provided at least 30 observations. Moreover, we need to take in account that four individuals of white-fronted lemur are related, what perhaps could have contributed to the dominance of right hand in this species, although the heritability of manual dominance is, at least, controversial, even inside one species. So, in chimpanzees, Hopkins (1999) did not find significant associations in hand preferences between parents and offspring, but in a further study, Hopkins et al. (2014) demonstrated that chimpanzee motor skill and handedness in tool use are significantly heritable.

Age and sex are factors that can affect the presence and direction of lateralization, as has been found in other studies with various species (McGrew & Marchant, 1997; Hopkins 2006; Schaafsma et al., 2008; Wells & Millsopp, 2009). Our results do not allow us to conclude anything clear about the effect of these factors on either species. The accentuation of behavioural lateralization with the age has been revealed in previous studies carried out on other primate species (Westergaard & Suomi 1993; Hopkins, 1995; Meunier et al., 2011), and especially on prosimians (Ward et al., 1990; Milliken et al., 1991; Shaw et al., 2004). As regard our study, in the white-fronted lemur they were observed in 6 adults and 1 juvenile, while in the ring-tailed lemur significant differences were observed in 2 adults and 1 juvenile. In neither species significant differences were observed in infants. In any case, it should be borne in mind that the sample in both species was clearly dominated by adult individuals.

On the other hand, although some authors claim that the sex of individuals may be a key factor determining the direction of lateralization (Milliken et al., 1989; Ward et al., 1990; Shaw et al., 2004) we cannot claim such an influence on the basis of our results. We cannot conclude anything about it in white-fronted lemur, and with respect to ring-tailed lemur, our results are somehow similar to those recorded by Schoeck & Edds (2018) in four individuals of this species: they found that the male was left-handed for feeding and the female was right-handed, while the rest, two twins, did not show significant differences. As far as our case is concerned, in white-fronted lemur significant differences were found in 4 males (2 left-biased, 2 right-biased) and 3 females (all right-biased), while in ring-tailed lemur, significant differences were observed only in 1 male (left-biased) and 2 females (1 left-biased, 1 right-biased). In neither species data are not sufficient to draw clear conclusions about the role of sex in the presence and the direction of lateralization.

## Conclusions

From the results obtained in this study we can conclude the following:

1. Feeding lateralization was more accentuated in white-fronted lemur than in ring-tailed lemur. In the first species a light bias towards the use of the right hand seems to be evidenced, although limitations of the samples must be taken into account.
2. From our results nothing can be concluded about the possible effect of age and sex on the presence and direction of lateralization.
3. The results do not seem to be in accordance with the postural origin of lateralization, since the more arboreal white fronted lemur was the species more lateralized concerning feeding behaviour, and the only that showed a light bias.

## Declarations

### Authorship clarified

Both authors contributed substantially to the elaboration of this work and approved the version to be published.

### Ethics approval

No approval of research ethics committees was required to accomplish the goals of this study because it was merely observational.

### Data Availability Statement

Data available on request from the authors.

